# Mast cell-derived chymases are essential for the resolution of inflammatory pain in mice

**DOI:** 10.1101/2024.08.05.606617

**Authors:** Sabrina de Souza, Sophie Laumet, Kufreobong E. Inyang, Hannah Hua, Jaewon Sim, Joseph K. Folger, Adam J. Moeser, Geoffroy Laumet

## Abstract

Immune cells play a critical role in the transition from acute to chronic pain. However, the role of mast cells in pain remains under-investigated. Here, we demonstrated that the resolution of inflammatory pain is markedly delayed in mast-cell-deficient mice. In response to Complete Freund Adjuvant (CFA), mast-cell-deficient mice showed greater levels of nitric oxide and altered cytokine/chemokine profile in inflamed skin in both sexes. In Wild-Type (WT) mice, the number of mast cell and mast cell-derived chymases; chymase 1 (CMA1) and mast cell protease 4 (MCPT4) increased in the inflamed skin. Inhibiting chymase enzymatic activity delayed the resolution of inflammatory pain. Consistently, local pharmacological administration of recombinant CMA1 and MCPT4 promoted the resolution of pain hypersensitivity and attenuated the upregulation of cytokines and chemokines under inflammation. We identified CCL9 as a target of MCPT4. Inhibition of CCL9 promoted recruitment of CD206^+^ myeloid cells and alleviated inflammatory pain. Our work reveals a new role of mast cell-derived chymases in preventing the transition from acute to chronic pain and suggests new therapeutic avenues for the treatment of inflammatory pain.

**Summary:** Mast cell-derived chymases play an unexpected role in the resolution of inflammatory pain and regulate the immune response.

**Graphical abstract:** Mast cells derived chymase MCPT4 degrades CCL9 to promote acute inflammatory pain resolution and prevent chronic pain. CFA-induced inflammation increases mast cells that degranulate and release chymases, like MCPT4 and CMA1, which in turn cleaves cytokines and chemokines such as CCL9. CCL9 cleavage induces the recruitment of CD206^+^ myeloid cells to promote the resolution of pain and prevent the transition from acute to chronic pain.

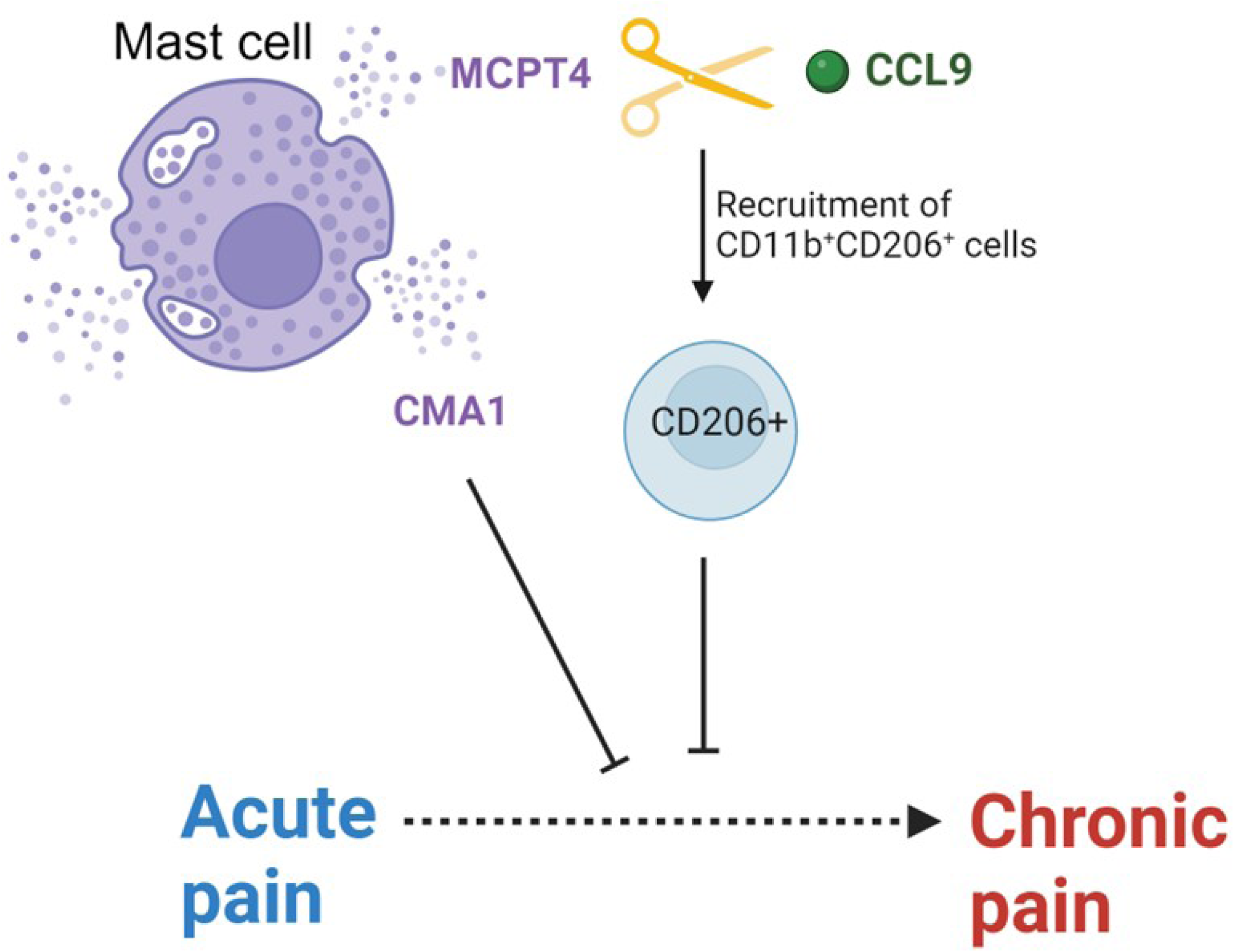

## Introduction

In the United States, chronic pain is a prevalent condition, impacting approximately 50.2 million adults, or 20.5% of the adult population [87]. It significantly impairs quality of life and worsens mortality outcomes compared to pain-free individuals [20,67,68].

It is well established that immune cells contribute to the development of chronic pain. Immune cells interact with neurons and modulate neuronal activity either activating or regulating the release of pro-inflammatory mediators (e.g., cytokines, chemokines, growth factors) that induce pain [15,76]. However, while the initiation of inflammatory pain is relatively well understood [12,37,80,86], the mechanisms behind the resolution and the contribution of specific immune cells remains unclear.

Mast cells reside in connective tissues and the skin and participate in the immune response and homeostasis [7,39]. Therefore, a tight interplay between nociceptors and mast cells has long been proposed based on anatomic colocalization [32,57,58]. Mast cells express high levels of c-Kit (CD117 or SCF receptor) and high-affinity IgE receptor (FceRI) [26]. When activated, they produce inflammatory mediators (e.g., TNF-α, IL-6), growth factors (e.g., TGF-ß1, platelet derived growth factor - PDGF) and proteases (e.g., tryptase and chymases). Proteases are the most abundant class of proteins produced uniquely by mast cells. Whereas tryptase is pro-inflammatory and mediates allergic inflammation, chymase has a regulatory role and contributes to tissue remodeling and fibrosis [9,17]. Although some chymases indeed increase in allergic inflammation (e.g. Mast cell protease 4), they are also protective and anti-inflammatory during infections [46,65,85,85]. Mast cell-derived chymases degrade pro-inflammatory cytokines to resolve inflammation [9,64,65,77,90] but their role in pain resolution has never been investigated.

While inhibition of mast cell degranulation relieves pain acutely [56], chronic activation of mast cells produces anti-inflammatory responses, increases interleukin (IL)-10 production and induces T regulatory cells (Treg) and Th2 T cells to promote peripheral tolerance [49]. In models of skin damage, mast cells have anti-inflammatory functions and prevent excessive swell, epidermis hyperplasia and necrosis, and leukocyte infiltration [33]. The long-term consequence of lack of mast cells in inflammatory pain has not been investigated.

Therefore, to investigate the role of mast cells in the transition from acute to chronic pain, we compared wild-type (WT) and mast-cell-deficient mice in response to a well-established model of inflammatory pain induced by the injection of Complete Freund Adjuvant (CFA). We found that although physiological nociception is similar in WT and mast-cell-deficient mice, inflammatory pain was significantly prolonged in the absence of mast cells. Manipulation of mast cell-derived chymases impacted cytokine and chemokine profile, such as CCL9 degradation, to mediate the resolution of inflammatory pain.

## Methods

### Mice and treatments

All animal experiments were conformed to the US National Institute of Health (NIH) guidelines and approved by Michigan State University (MSU) Institutional Animal Care Use Committee (IACUC; protocol #201900249-202200290). C57BL/6 (JAX#000664, WT) and Kit^W-sh/W-sh “Sash” mice (JAX#030764) were purchased from Jackson laboratory (Bar Harbor, ME) and housed and bred in the MSU Animal Care Facility. Equal number of 8-10 weeks old female and male mice were used in experiments.

Complete Freund’s adjuvant (CFA; Sigma-Aldrich F5881) was injected unilaterally into the plantar surface of the hind paw in WT and Sash mice (5 ul) under isoflurane anesthesia. Chymostatin (Sigma-Aldrich; 230790-10MG) was intraperitoneally administrated. Recombinant CMA1 (Sigma-Aldrich; C8118), recombinant MCPT4 (MyBioSource; MBS1133087), or anti-CCL9 antibody (Peprotech; 500-P117) was administrated into the hind paw.

### Pain sensitivity assessment Heat pain

Mice were placed in a 52°C metal surface after adaptation at 26°C, and the latency to a nociceptive response was recorded, defined as the time elapsed until the mice lick or flicks its hind paw and/or jumps.

### Mechanical pain

Mechanical sensitivity was assessed using von Frey filaments (0.04-1.4 g) as a response to physical stimulation on the plantar surface of the hind paw. Briefly, after habituation, von Frey filaments were applied on the plantar surface, increasing forces (grams) as previously described (Inyang, 2023). For response frequency, each filament was used 10 times. Mice were considered recovered upon reaching 80% of their baseline values.

### Cold pain

Cold pain was measured by spraying 0.1 ml of acetone to the plantar surface of the hind paw, eliciting cooling of the skin and the responsive time was recorded, defined as time the mice look and interact to the hind paw.

### Skin collection

Mice were euthanized using carbon dioxide and hind paw skin were harvested either for staining, RNA isolation using Trizol chloroform method (Heussner et al., 2021) (Ambion Life technologies; 15596018), processed using lysis buffer (Ray Biotech; EL-lysis) with protease inhibitor cocktail (Sigma Aldrich; 118361530001) for protein assessment and cytokine array or processed for flow cytometry.

### Gene expression

Samples were sonicated (Fisher brand CL-18), RNA isolated and RNA concentration was measured (Fisher Scientific, Q33211) with a Qubit 4 fluorometer. Samples were stored at −80⁰C until further use. Gene expression was accessed by quantitative Real-Time polymerase chain reaction (qRT-PCR) after RNA isolation and cDNA synthesis by reverse transcription. High-Capacity cDNA Reverse Transcription Kit (Applied biosystem, Thermo Fisher Scientific; 4368814) was used for cDNA synthesis in T100TM Thermal Cycler (BioRad®) under the following conditions: 10 minutes at 25°C, 120 minutes at 37°C, 5 minutes at 85°C and samples were diluted and stored at −20°C until use. qRT-PCR were performed using CFX Connect™ Real-Time System (BioRad®) instrument and BioRad CFX Manager Software. qPCR was performed using validated PrimeTime primer assays for *Gapdh* (Ex2-3#Mm.PT.39a.1(1)), *Cma1* (Ex1-2#Mm.PT.584586134) and *Mcpt4* (Ex2-3#Mm.PT.58.32178595) (Integrated DNA technologies). Relative expressions of genes were normalized to the internal reference *Gapdh* and analyzed using 2−ΔΔCT method.

### Nitric oxide

Nitric oxide (NO) was quantified using Griess method (Molecular Probes; G-7921) according to manufacturer instructions. Colorimetric optical densities (OD) were measured at 560 nm by Infinite F50 (Tecan®), and the concentrations were analyzed using GainData Software (Arigo Biolaboratories, https://www.arigobio.com/elisa-analysis).

### Mast cell staining

Paw skin was collected from WT and Sash mice and stained with Toluidine blue (Sigma-Aldrich, 92-31-9). Briefly, after 15 minutes fixation with 3% glutaraldehyde in PBS, samples were rinsed with milli-Q water and permeabilized with 0.5% Triton X-100 for 5 minutes. Then, samples were rinsed again and stained with 0.3% Toluidine blue (prepared in Milli-Q water and filtered) for 30-60 seconds. Images were taken with Nikon Eclipse NI microscope.

### Protein Cytokine-array

Paw skins were processed with Lysis buffer (Ray Biotech; EL-LYSIS) for Mouse Cytokine Array 3 (Ray Biotech; AAM-CYT-3-4). After sonication and centrifugation (5,000 x g for 5 minutes), 2,000 ug/ml of sample were incubated on the membranes containing 62 murine cytokines specific antibodies, blocked and incubated for 5 hours at room temperature with gentle shaking. After washing, membranes were incubated with biotinylated antibody overnight at 4°C. Next day, membranes were washed and incubated with Streptavidin-HRP for 2 hours, washed once again and transferred into a sheet provided by Ray Biotech for 2 minutes incubation with the detection antibody. Membranes were scanned using Omega Lum G imaging. Density was extracted and background subtracted using ImageJ. Arbitrary units were obtained following instructions provided by the manufacturer. Relative expression was normalized using the formula Y=X-MIN(Row)/MAX(Row)-MIN(Row), considering 0 as minimum as 1 as maximum expression. Raw data represented on supplementary tables 1-3.

### Single cell suspensions and flow cytometry

Plantar skin tissues were collected and immediately placed in ice-cold 1X Gibco™ HBSS (HBSS (10X), 14-185-052, Waltham, MA). Two or three paw skins were pooled together to obtain > 20,000 immune cells. Mouse skin immune cells were isolated as previously described [13,74]. Tissues were diced into 1 mm-sized pieces to facilitate enzymatic digestion (5 ml of RPMI-1640 Medium - R7509, Sigma, St. Louis, MO; 1 mg/ml DNase I - DN25, Sigma-Aldrich, St. Louis, MO - and 1 mg/ml Collagenase D - 11088858001, Sigma, St. Louis, MO). Enzymatic digestion was performed at 37°C for 2 hours. Digested tissues were minced and filtered through a 70 μm cell strainer by grinding tissue gently with the plunger end of the 10 ml syringe. After centrifugation at 400 g for 10 minutes at 4°C, supernatants were discarded, and the pellets were resuspended with 2 ml of PBS for staining and flow cytometry.

Cells were first stained with 1 μL/mL of LIVE/DEAD Blue (L34962, Invitrogen, Carlsbad, CA) for 15 min on ice. Fc-blocking antibodies CD16/CD32 (553142, BD, Franklin Lakes, NJ) were added to single-cell suspension prior to surface staining. Cells were then stained with CD45-BUV563, CD11b-APC-fire-750, NK1.1-BUV805, CD3-APC-fire-810, Fcer1-Super Bright 600, and CD11c-BV650 in flow cytometry staining buffer (PBS + 1% Bovine Serum Albumin-BSA) and brilliant staining buffer (563794, BD, Franklin Lakes, NJ) at 4°C for 1 hour in the dark. Cells were analyzed with a 5 laser (16UV-16V-14B-10YG-8R) Cytek® Aurora - Spectral Flow Cytometry (Cytek Biosciences, Fremont, CA) located in the MSU Flow Cytometry Core Facility, and the data were unmixed with SpectroFlo® software and analyzed using FlowJo™ v10.8 Software (BD Life Sciences). Ultracomp eBeadsTM (01-2222-41, Invitrogen, Carlsbad, CA) stained with antibodies were used as the reference controls for unmixing. To prevent interference from the autofluorescence, unmixing algorithms in the SpectroFlo software were used to extract autofluorescence from unstained cells.

### Chymase activity

Chymase activity was determined in processed paw skin by the hydrolysis rate of 10 uM of the fluorogenic substrate, Suc-Leu-Leu-Val-Tyr-7-amino-4-methylcoumarin (MCE, HY-P1002) at 37°C for 1 hour. Fluorescence was measured on SpectraMax™ M5 spectrophotometer with λex=370 nm and λem=460 nm. The concentration (nM) of the fluorogenic substrate cleavage was determined based on a standard curve [41].

### Detection of CCL9

The concentration of MIP-1-γ/CCL9 (RayBiotech; ELM-MIP1g-1) in paw skin lysate (processed with Lysis buffer, Ray Biotech; EL-LYSIS) was quantified by ELISA according to manufacturer’s instructions. The concentrations were analyzed by GainData Software and normalized by protein.

### Statistical analysis

Experimental data were analyzed using GraphPad Prism 9.4.1 Software. Values expressed on graphs indicate the mean ± standard error (SEM) from independents experiments. Differences were analyzed by unpaired t test or two-way ANOVA with multiple comparison test, according to experimental design. Values of p < 0.05 were considered significant.

## Results

### Naïve mast cell-deficient and WT mice show similar pain sensitivity

The c-kit tyrosine kinase is essential for mast cell survival [35], therefore mice carrying mutations in the white spotting locus of *Kit* gene, so-called Sash mice, are virtually deficient of mast cells [28]. Toluidine blue staining confirmed the absence of mast cells in the skin and qPCR revealed a marked reduction of *Fcer1* expression (**Figure 1A-B**). In naïve conditions, WT and mast cells deficient (Sash) mice have similar response to heat (**Figure 1C**), cold (**Figure 1D**) or mechanical (**Figure 1E**) stimuli, showing that the deficiency of mast cells does not affect nociception in the absence of inflammation.

**Figure 1:**
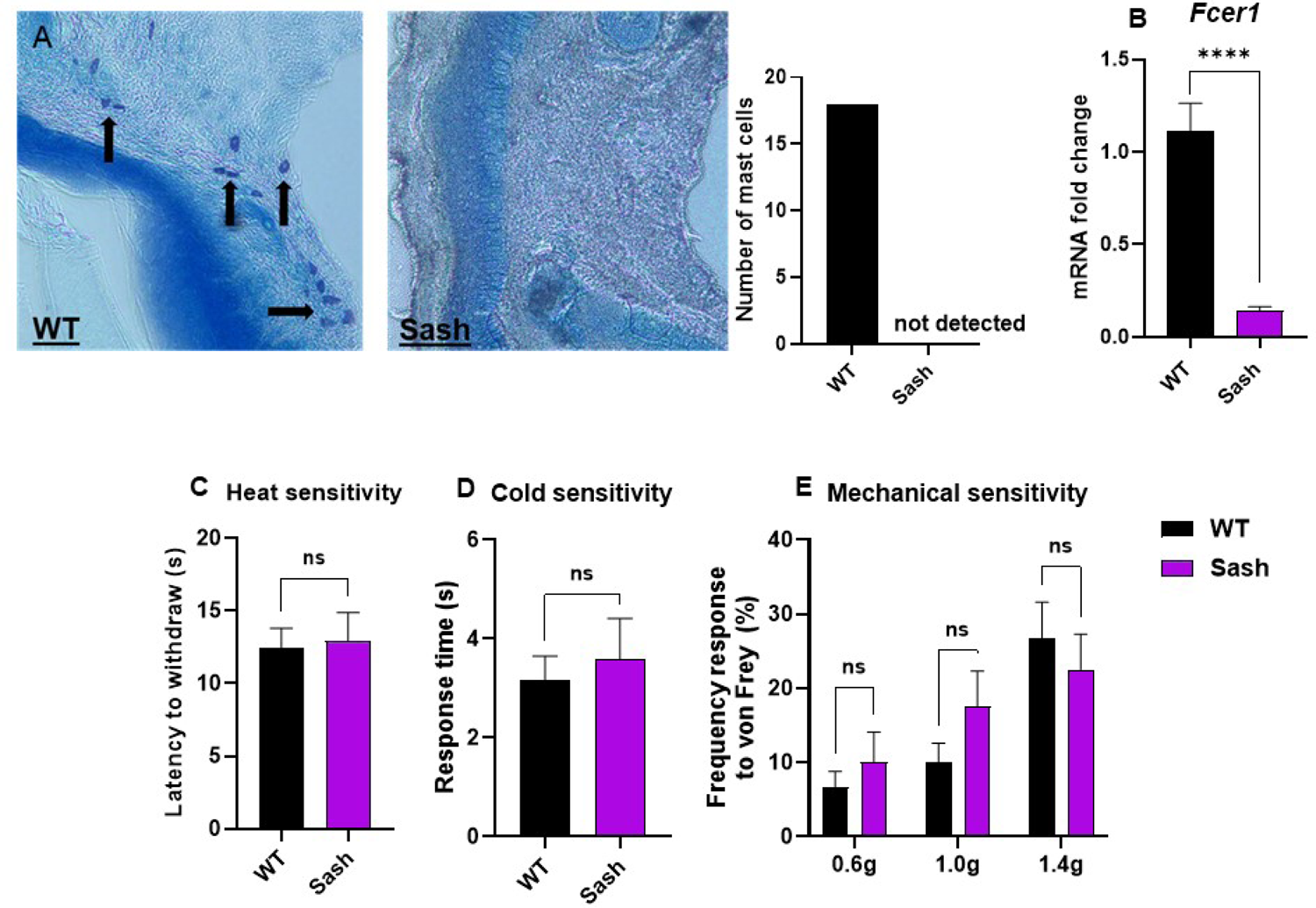
Pain sensitivity is preserved in mast cell-deficient naïve mice. A-B) Mast cells deficiency validation. A) Plantar paw skin was collected from WT and Sash mice and prepared for toluidine blue coloration. Left panel shows the presence of mast cells in WT (arrows) and right panel shows the absence of mast cells in Sash. Images taken with fluorescent microscope at 20x objective. Graph shows the quantification of mast cells in WT and Sash. B) *Fcer1* mRNA fold change expression in paws skin of naïve WT and Sash mice. Two-tailed Mann-Whitney test; p<0.0001. C) Withdrawal latencies of WT (n=7) and Sash (n=7) mice on the hot plate at 52°C. Two-tailed unpaired test p=0.8539. D) Response time of WT and Sash mice to acetone. Two-tailed unpaired test; p=0.676. E) Response frequency of WT (n=6) and Sash (n=4) mice against von Frey filaments (0.6, 1.0 and 1.4g). Two-tailed unpaired test p=0.4468 0.6 g; p=0.1700 1.0 g, p=0.5811 1.4 g. Results are presented as mean ± SEM from independent experiments.

### The number of mast cells increased in inflamed skin

To explore the impact of mast cells on inflammatory pain, we injected CFA in the hind paw of WT mice and performed flow cytometry for mast cells identification. Mast cells were determined as CD45^+^ CD11b^-^ CD11c^-^ CD3^-^ NK1.1^-^ Fcer1^+^ (**Figure 2A**). Seven days after administration of CFA induced a significant accumulation of immune cells (CD45^+^, **Figure 2B**). Interestingly, the number of mast cells significantly increased in the inflamed skin (**Figure 2C**).

**Figure 2:**
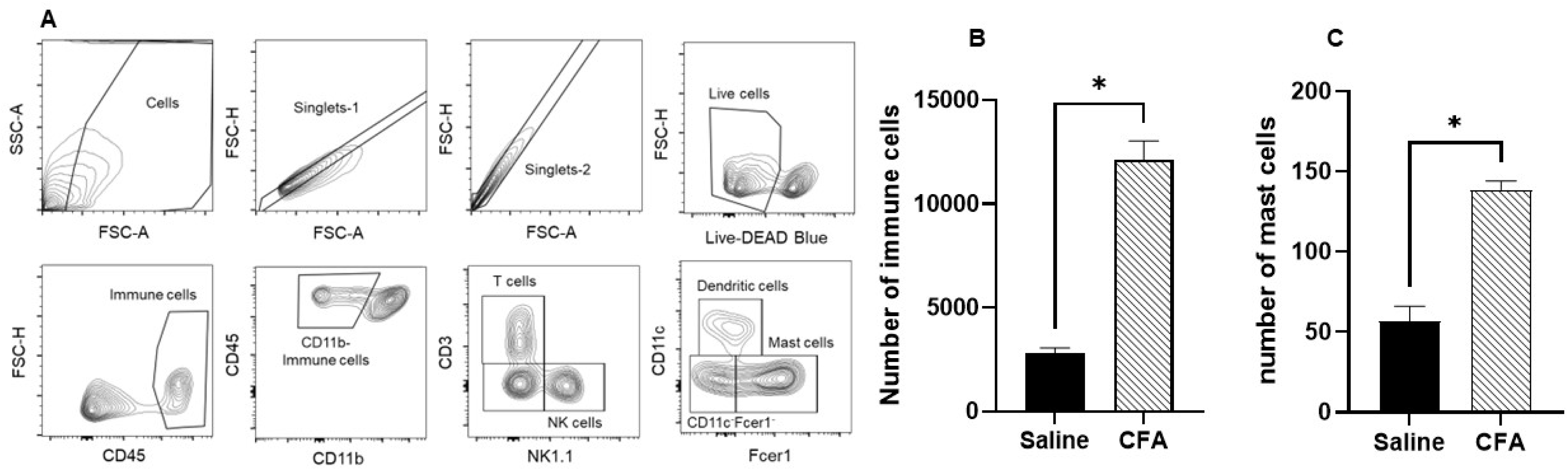
CFA increases the number of immune cells and mast cells in the inflamed skin. A) Representative flow plots of gating strategy used for immune cell identification based on forward scatter (FSC-A) and side scatter (SSC-A), singlets, live cells and CD45^+^ cells. Immune cells (CD45^+^) were gating into population according to CD11b, NK1.1, CD3, CD11c and FCERI expression. Mast cells were characterized as CD11b^-^ CD11c^-^ FCERI^+^. B) Absolute number of immune cells 7 days after saline (control) and CFA. Two-way ANOVA test with multiple comparisons; p= 0.0186. C) Absolute number of mast cells after treatments. Two-way ANOVA test with multiple comparisons; p=0.0286 (C). Mean ± SEM from 3 (saline) and 4 (CFA) independent experiments.

### Mast cell deficiency impacts the resolution of pain hypersensitivity induced by CFA

Given that mast cells increased in the inflamed skin, we next evaluated the effect of mast cells on CFA-induced pain hypersensitivity. CFA induced robust mechanical hypersensitivity, demonstrated by decreased mechanical threshold of both WT and Sash after 24 hours (**Figure 3A**). However, whereas WT recovered in 12 days, mast cell-deficient mice did not recover within the testing period (**Figure 3A-B, Two-way ANOVA genotype effect F (1,98) = 63.88, p<0.0001**), suggesting a beneficial role of mast cells in the resolution of inflammatory pain. CFA-induced edema was more pronounced in Sash mice compared to WT (**Figure 3C, Two-way ANOVA genotype effect F (1,14) =10.5, p<0.006**), implying potentially more intense inflammation in the absence of mast cells. To confirm this idea, we measured the levels of nitric oxide (NO) as a marker of inflammation. Sash mice had a greater increase of NO in the inflamed skin compared to WT mice (**Figure 3D**). Using a cytokine and chemokine array, we identified more than 30 cytokine/chemokine proteins differentially regulated in the inflamed skin of WT and Sash mice 8 days after CFA (**Figure 3E, Two-way ANOVA, genotype effect F (2, 118) = 34.66, p<0.0001; cytokine effect F (59, 118) = 2,757, p<0.0001; supplementary table 1**).

**Figure 3:**
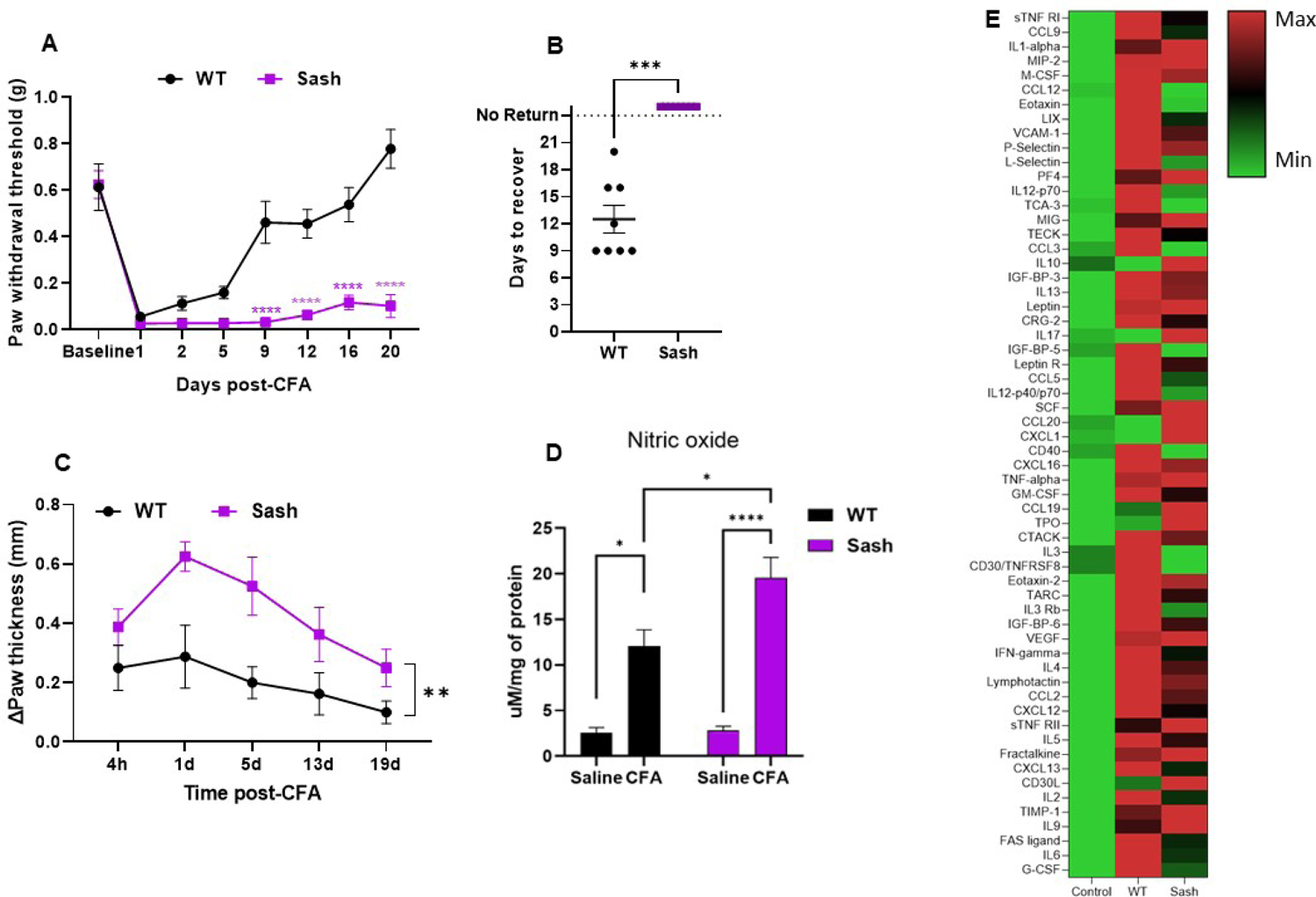
Absence of mast cells results in greater inflammation and impaired the resolution of inflammatory pain. WT and Sash mice were subjected to plantar injection of CFA after baseline determination. A) Paw mechanical pain sensitivity in response to von Frey filaments after CFA injection (n=8). Two-way ANOVA with multiple comparison; p<0.0001 compared to its control. B) Time to recover from CFA-induced inflammation. Two-tailed Mann-Whitney t test; p=0.0002. C) Paw thickness of injected paw (n=8). Two-tailed Mann-Whitney t test. D) Levels of nitric oxide in saline (control) and CFA-injected paws, normalized by protein concentration (mg/ml). Two-way ANOVA with multiple comparison; *p=0.0161, ****p<0.0001. E) Heatmap representing magnitude of chemokine and cytokine protein released in WT (pool of 3 male + 3 female) and Sash (pool of 3 male and 3 female) mice 8 days after CFA.

### Mast cells chymases mediate pain resolution

Chymases are the major proteases secreted uniquely by mast cells [3]. Mouse protease 4 (MCPT4) and chymase 1 (CMA1 or MCPT5) are predominantly expressed in skin-resident mast cells [6]. These proteases have been associated with relieving of inflammation by degrading inflammatory mediators such as TNF-α [65,90] and chemokines MIP-3a/CCL20 and MIP-3b/CCL19 [22].

The expression of *Mcpt4* and *Cma1* were upregulated in CFA-treated paws (**Figure 4A-B**). Chymase enzymatic activity also significantly increased 8 days after CFA, while chymase activity **was absent in Sash mice (**Figure 4C-D, Two-way ANOVA, treatment effect F (1, 84) = 122.7, p<0.0001; time effect F (11, 84) = 103.8, p<0.0001).

**Figure 4:**
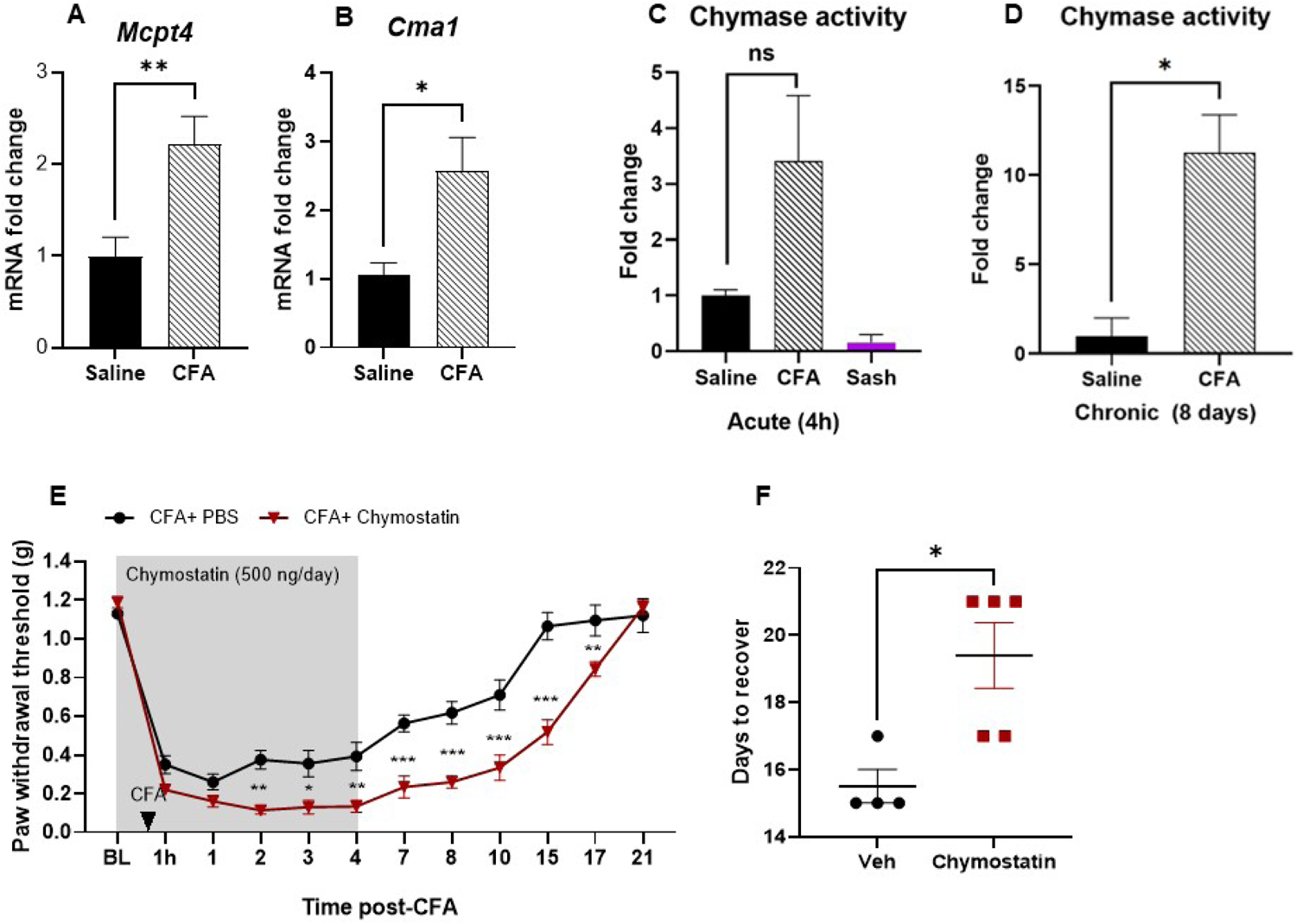
Chymase activity contributes to the resolution of inflammatory pain hypersensitivity. A-B) *Mcpt4* and *Cma1* mRNA levels in saline (control) and CFA-treated paw skin. Two-tailed Mann-Whitney t test; *p=0.0104 (A), **p= 0.0059 (B). C-D) Chymase activity at 4h (C, acute phase) and 8 days (D, chronic phase) after CFA-induced inflammation, measured as AMC-specific cleavage of the fluorogenic substrate Suc-Leu-Leu-Val-Tyr-AMC. One-way ANOVA test with multiple comparisons; p=0.5700 (C); t-tailed Mann-Whitney t test; p= 0.0303 (D). E) Paw withdrawal threshold accessed by von Frey filaments after CFA and PBS (control) (n=5) or chymostatin (500 ng/day) injection (n=4). Two-tailed Mann-Whitney t test, *p=0.0263, **p=0.0052, ***p=0.0002, ****P<0.0001. F) Time to recover from CFA + vehicle or chymostatin injection. Two-tailed Mann-Whitney t test; p= 0.0159. Results are presented as mean ± SEM from independent experiments.

To gain insight into the contribution of mast cell-derived chymases to inflammatory pain, we systemically inhibited chymase enzymatic activity using the specific pharmacological inhibitor chymostatin [34]. Administration of chymostatin significantly delayed the resolution of pain hypersensitivity (**Figure 4E, Two-way ANOVA, treatment effect F (1, 84) = 122.7, p<0.0001; time effect F (11, 84) = 103.8, p<0.0001**) and prolonged the return to baseline by 5 days (**Figure 4F**).

Given that inhibition of chymase activity impaired the resolution of pain hypersensitivity, we next determined whether injection of MCPT4 or CMA1 facilitates the resolution of pain hypersensitivity. Administration of recombinant MCPT4 or CMA1 in the inflamed skin alleviated inflammatory pain in WT mice (**Figure 5A-B**) and significantly fastened the return to baseline (**Figure 5C**). In addition to alleviating pain hypersensitivity, administration of MCPT4 reversed the upregulation of cytokine and chemokine proteins in response to CFA (**Figure 5D-E; supplementary table 2**).

**Figure 5:**
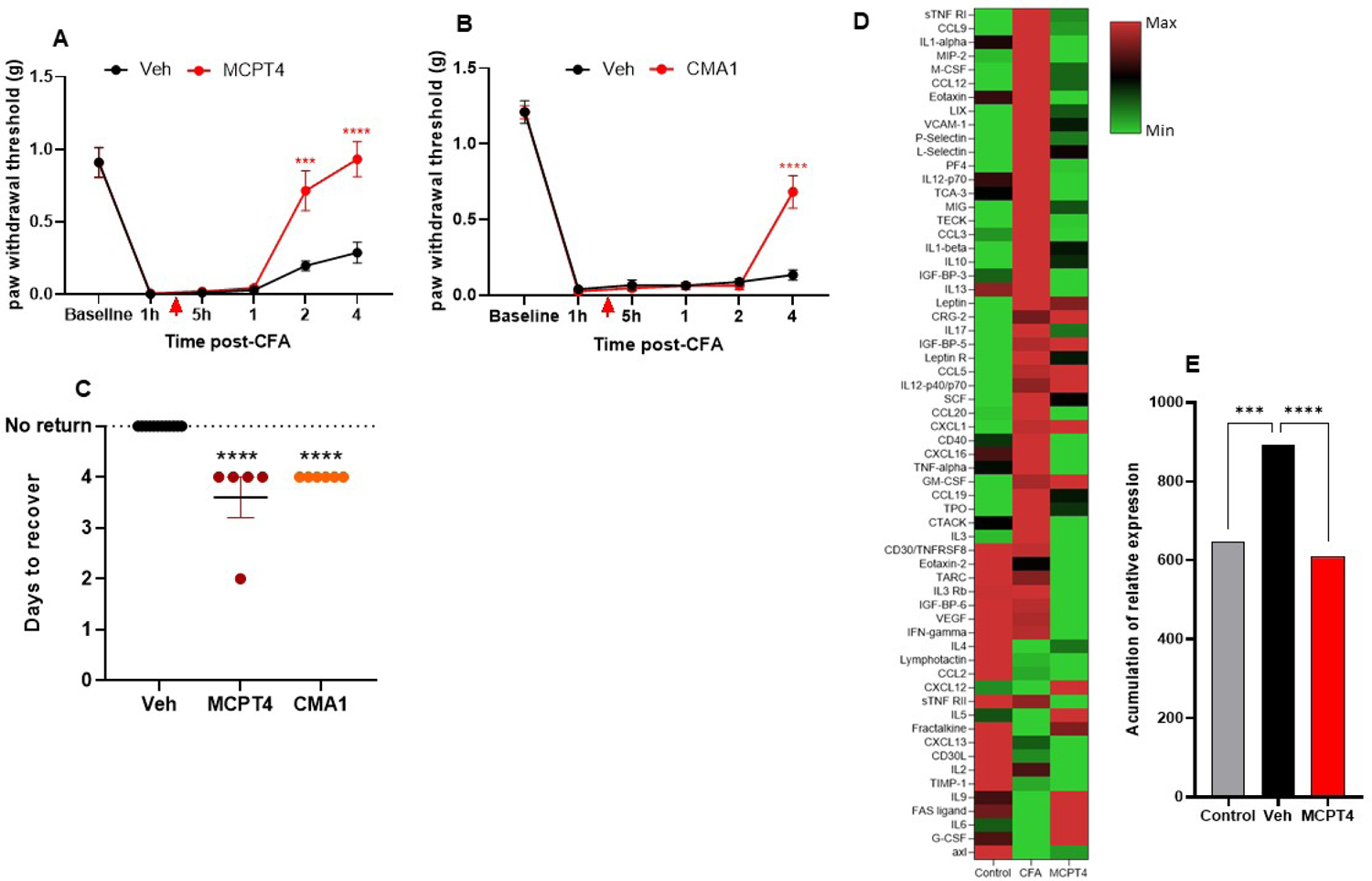
Administration of mast cell chymase MCPT4 and CMA1 ameliorates pain resolution in WT mice: A) Paw withdrawal threshold of WT mice in response to von Frey filaments after vehicle (PBS control) or 1ug of recombinant MCPT4 (A) (n=5) administrated 4 hours after CFA injection (red arrow). Two-way ANOVA test with multiple comparisons; ***p=0.0003, ****p<0.0001. B) Paw withdrawal threshold of WT mice after vehicle (PBS) and CMA1 (n=8) administrated 4 hours after CFA. Two-way ANOVA test with multiple comparisons; ****p<0.0001. C) Time to recover from CFA + vehicle, CFA + MCPT4 or CMA1. Ordinary one-way ANOVA; p<0.0001. D) Heatmap representing magnitude of cytokines and chemokines in control (naive paw), CFA + vehicle and CFA + MCPT4. Pool of 3 males and 3 females/condition. E) Accumulation of relative expressions in control, CFA + vehicle and CFA + MCPT4. Two-way ANOVA with multiple comparisons; ***p= 0.0004, ****p<0.0001.

To determine whether injection of recombinant chymases compensate for the lack of mast cells, we administered recombinant MCPT4 or CMA1 in the inflamed skin of Sash mice, but neither of the chymases affected pain hypersensitivity individually (data not shown). Strikingly, combination of MCPT4 and CMA1 significantly alleviated pain hypersensitivity and triggered recovery in Sash mice (**Figure 6A-B)**. In Sash mice, MCPT4 and CMA1 also reversed exaggerated inflammatory cytokines and chemokines (**Figure 6C-D, two-way ANOVA genotype effect F (2,100) = 31.04, p<0.0001; cytokine effect F (50,100) = 2693, p<0.0001; supplementary table 3**).

**Figure 6:**
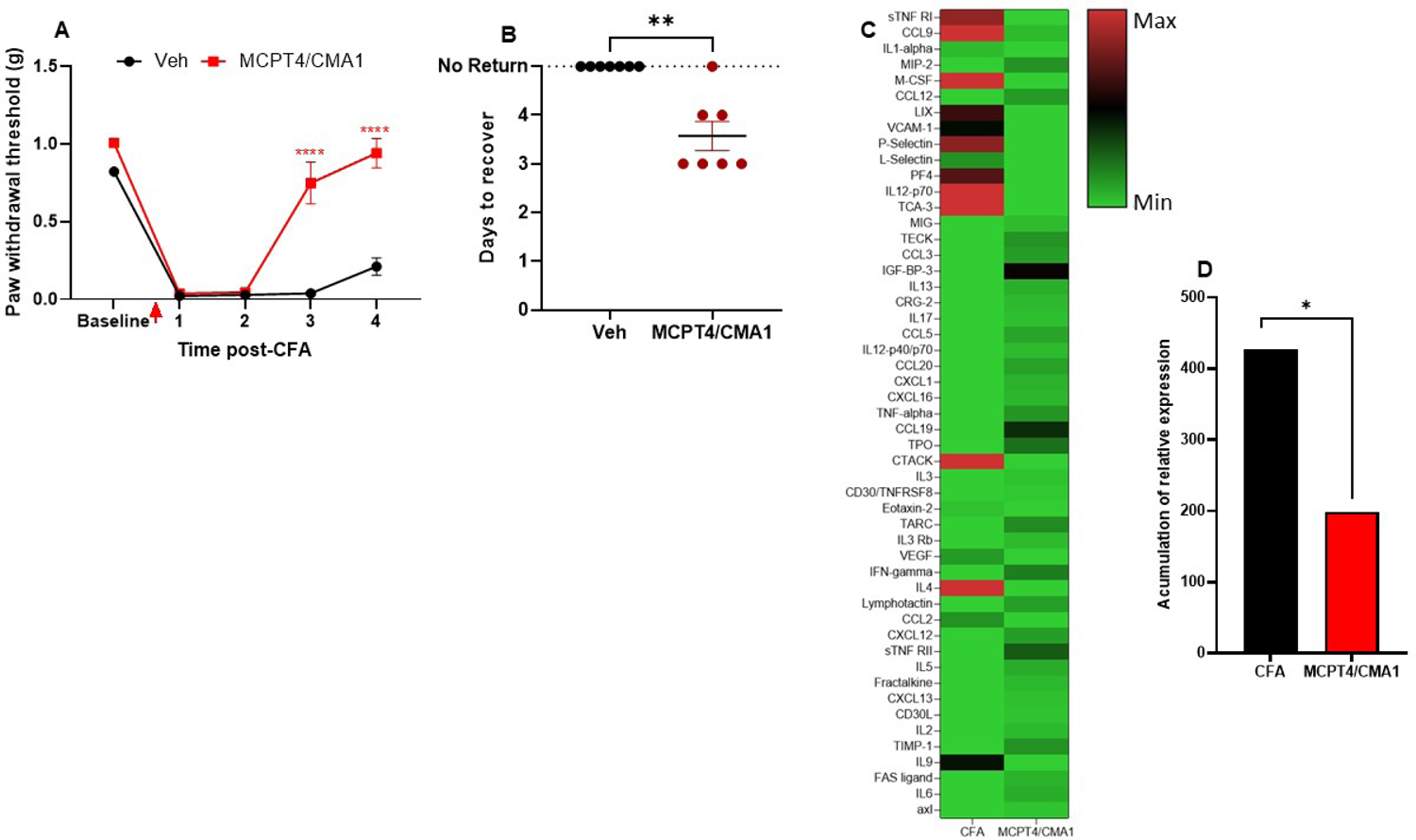
MCPT4 and CMA1 improves pain in mast cell deficient mice: A) Paw withdrawal threshold of Sash mice after CFA and MCPT4 combined with CMA1. Two-way ANOVA test with multiple comparisons; p<0.0001. B) Time to recover from CFA + vehicle and CFA + MCPT4/CMA1. Two-tailed Mann-Whitney t test; p= 0.0006. C) Heatmap representing magnitude of cytokines and chemokines in WT + CFA + vehicle and Sash mice injected with CFA + MCPT4/CMA1. Pool of 3 males and 3 females/condition. D) Accumulation of relative expressions. Two-way ANOVA; *p= 0.0127.

### MCPT4 degrades CCL9 to promote the resolution of pain

Chemokine arrays revealed that CCL9 is reduced by addition of MCPT4 in both WT and Sash mice. CCL9 is degraded by mast cell protease under inflammatory conditions [5]. Therefore, we investigated whether CCL9 regulation by MCPT4 mediates the resolution of pain. ELISA experiment indicated that levels of CCL9 significantly increased in CFA-treated sample and reduced after administration of MCPT4, strongly suggesting that CCL9 may be degraded by MCPT4 (**Figure 7A**).

**Figure 7:**
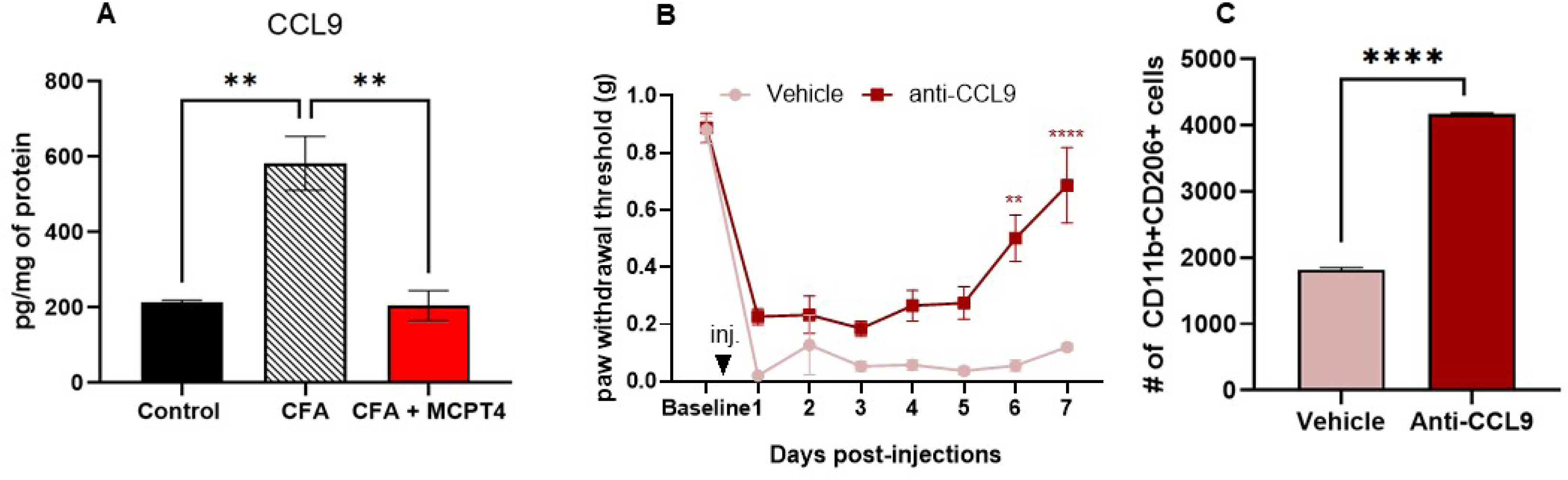
MCPT4 regulates chemokine CCL9 levels to promote pain resolution via CD206+ myeloid cells. A) Protein levels of CCL9 assessed by ELISA in paw skin in control (naive paw) (n=5), CFA + vehicle (n=9) and CFA + MCPT4 (n=5), 4 days after injections. Ordinary one-way ANOVA with multiple comparisons; p=0.0020. B) Paw withdrawal threshold of WT mice treated with CFA + vehicle or anti-CCL9 (injected 4 hours after CFA). Two-way ANOVA test with multiple comparisons; **p=0.0010; ****p<0.0001. C) Absolute number of CD11b^+^ CD206^+^ cells in CFA + vehicle and CFA + anti-CCL9 treated skin (4 hours after CFA). Samples collected on day 5 post-injections. Chi-square with Fisher’s exact test. ****p<0.0001.

CCL9 has been reported to have pronociceptive properties [63,69] and pharmacological manipulation of its receptor, CCR1, has been shown to reduce neuropathic pain [14]. Here, injection of CCL9 neutralizing antibody reduced CFA-induced pain hypersensitivity and triggered the resolution of pain (**Figure 7B**).

Studies have shown that CCL9 receptor is expressed by CD206+ macrophages and regulate their activity [16,43,54,89]. This is interesting because anti-inflammatory immune cells, such as CD206^+^ macrophages are essential for the resolution of pain [12,25,60,74,75,83]. Therefore, we evaluated if CD206+ cells recruitment is regulated by CCL9. The neutralization of CCL9 increased the number of CD206^+^ myeloid cells (likely to be anti-inflammatory macrophages) (**Figure 7C**) in the inflamed skin following CFA injection. These results suggest that the cleavage of CCL9 by MCPT4 contributes to the resolution of inflammatory pain by recruiting anti-inflammatory CD206^+^ macrophages.

Altogether, our findings indicate that mast cell-derived chymases, MCPT4 and CMA1, are necessary for the resolution of CFA-induced inflammatory pain by degrading cytokines and chemokines, such as CCL9. CCL9 controls the recruitment of CD11b^+^ CD206^+^ immune cells, which help to resolve acute inflammatory pain and, therefore, prevent the transition from acute to chronic pain (**Figure 8**).

## Discussion

Although immune cells contribute to the development of chronic pain, an emerging body of studies have shown a critical role of immune cells in the resolution of pain [4,42,74,83]. Here, we showed, for the first time, that mast cells and more specifically mast cell-derived proteases, like MCPT4 and CMA1, are essential to prevent the transition from acute to chronic pain following inflammation. We further showed that MCPT4 cleaves the chemokine CCL9, facilitating the recruitment of CD11b^+^CD206^+^ myeloid cells to mediate the resolution of inflammatory pain.

Proteases are the most abundant class of proteins produced uniquely by mast cells, which mainly include chymases and tryptases [1,8,64]. Studies have shown the role of tryptases inducing pain hypersensitivity via PAR2 [21,40,52,55,61,70,82]. While mast cell tryptases have been often investigated in pain models, the role of chymases in pain is unknown. Chymases degrade various proteins such as neuropeptides substance P (SP), calcitonin gene-related peptide (CGRP) and vasoactive intestinal peptide (VIP) [10,79], as well as pro-inflammatory mediators HMGB1, bradykinin, IL-6, TNF-α and IL-33 [36,65,71,77,85,90]. These proinflammatory mediators are known to facilitate nociception [66,73]. Previous studies demonstrated that mast cell chymases are protective in the context of neuroinflammation. Administration of MCPT6 promotes recovery from spinal cord injury and reduction of TNF-α and IL-1b [81]. MCPT4 was also protective in models of traumatic spinal cord and brain injury, by degrading pro-inflammatory cytokines [33,59]. MCPT4 also has a protective role in allergic inflammation [84,85] and maintains barrier homeostasis [30,51]. In the present study, we speculate that CMA1 and MCPT4 degrade proinflammatory/pronociceptive inflammatory mediators to facilitate the resolution of inflammation and pain. Indeed, administration of MCPT4 reduced the protein levels of several pro-inflammatory cytokines. Specifically, we showed that MCPT4 degrades CCL9, a chemokine that regulates the recruitment of myeloid immune cells [54,90] and it is degraded by mast cells chymases under inflammatory conditions [5].

We do not claim that CCL9 degradation is solely responsible for MCPT4’s effects on the resolution of pain. Although neutralizing CCL9 was sufficient to promote the resolution of pain, it is likely that the degradation of other cytokines and chemokines by chymases contributes to the transition from acute to chronic pain. It is also possible that chymases resolve nociception through additional mechanisms, such as direct action on nociceptors. These possibilities will be explored in future studies.

CCL9 has gained interest in cancer immunology because of its ability to modulate tumor-associated macrophages, which are CD206^+^ myeloid cells. Recently, CD206^+^ myeloid cells have attracted a lot of attention in pain research because of their capacity to promote the resolution of pain [4,25,60,74,75,83]. Initial activation of mast cell-derived chymases might be necessary for the recruitment of CD206^+^ myeloid cells and the prevention of the transition to chronic pain.

The contribution of mast cells to pain is controversial. Histamine and serotonin released from mast cells contribute to neuronal sensitization [31,45,78,88]. Acute inhibition of mast cell degranulation improves hyperalgesia [2,53,56,82]. Mast cell stabilizer ketotifen fumarate reverses inflammatory pain [56] but long-term consequences of mast cell inhibition have never been investigated. Here we took advantage of genetic model to study the long-term depletion of mast cells on inflammatory pain. The onset and severity of mechanical hypersensitivity were similar in WT and Sash mice, but the resolution of pain was markedly delayed in absence of mast cells. Indeed, mast cells are known to limit the immune response [23,29,44,65,71]. While acute stimulation triggers degranulation and release of proinflammatory molecules, chronic stimulation produces anti-inflammatory response facilitating the recruitment of T regulatory cells (Treg), Th2 T cells, and IL-10-producing cells [29,48,49]. Specifically in the skin, in models of ultraviolet radiation-induced skin damage, mast cells have anti-inflammatory functions and prevent excessive leukocyte infiltration [29]. Additionally, mast cells promote the resolution of contact hypersensitivity and inflammation in response to poison ivy [29]. They release IL-4, IL-10 and TGF-β that reduce pain and inflammation [18,47,72].

Interestingly, our cytokine/chemokine protein arrays showed that the absence of mast cells drastically alters the immune response. We did not only observe an increase in expression, but some cytokines were higher in Sash compared to WT and others were lower in Sash compared to WT indicating an important role of mast cells in the regulation of skin inflammation.

Based on the inflammatory profile, CCL9 appears to be interesting candidates as causing nociception and being degraded by mast cell-derived chymases. Indeed, both cytokines are upregulated by CFA, higher in Sash mice, and reduced by administration of recombinant chymases in both WT and Sash. We confirmed the degradation of CCL9 by MCPT4 which resulted in the recruitment of CD206^+^ cells. Establishing the potential implication of CCL9 on inflammatory pain will require further investigation.

Although the immune system is sexually dimorphic [19,24,38] and neuro-immune interactions have been suggested to contribute to sex differences in pain [27,50,62,74], here we did not observe any sex differences. In the present study, the absence of mast cells drastically prolonged the resolution of pain in both sexes, upregulation of chymases was similar in both sexes, and exogenous administration of recombinant chymases promote the resolution in both sexes.

Our findings showed that mast cell-derived MCPT4 and CMA1 are essential for the natural resolution of inflammatory pain and preventing the acute-to-chronic pain transition. But we also showed that administration of MCPT4 and CMA1 accelerated the resolution of inflammatory pain in WT and in absence of mast cells. These chymases are both necessary and sufficient to prevent the transition into chronic pain.

In the present study, we identified mast cell-derived chymases as critical endogenous “brake” for the acute-to-chronic pain transition. Future studies will characterize the precise molecular underlying the resolution of pain driven by chymases. Our study reveals administration/activation of mast cell-derived chymases and neutralization of CCL9 as new therapeutic strategies to prevent chronic pain.

## Conflict of Interest statement

None of the authors have any conflicts of interest to declare.

## Acknowledgement

This work was supported by the Rita Allen Foundation, National Institute of Health (NIH R01NS121259), the Department of Defense (CPMRP CP220093) to G.L, and the NIH (R01 HD072968, R01AI168014) to AJM.

## Data availability statement

Original data will be made available upon reasonable request.

## Notes

### Competing Interest Statement

The authors have declared no competing interest.

